# Dopamine D1-like receptors modulate synchronized oscillations in the hippocampal-prefrontal-amygdala circuit in contextual fear

**DOI:** 10.1101/2023.06.15.545080

**Authors:** Christine Stubbendorff, Ed Hale, Tobias Bast, Helen J Cassaday, Stephen J Martin, Sopapun Suwansawang, David M Halliday, Carl W Stevenson

## Abstract

Contextual fear conditioning (CFC) is mediated by a neural circuit that includes the hippocampus, prefrontal cortex, and amygdala, but the neurophysiological mechanisms underlying the regulation of CFC by neuromodulators remain unclear. Dopamine D1-like receptors (D1Rs) in this circuit regulate CFC and local synaptic plasticity, which is facilitated by synchronized oscillations between these areas. In rats, we determined the effects of systemic D1R blockade on CFC and oscillatory synchrony between dorsal hippocampus (DH), prelimbic (PL) cortex, basolateral amygdala (BLA), and ventral hippocampus (VH), which sends hippocampal projections to PL and BLA. D1R blockade altered DH-VH and reduced VH-PL and VH-BLA synchrony during CFC, as inferred from theta and gamma coherence and theta-gamma coupling. D1R blockade also impaired CFC, as indicated by decreased freezing at retrieval, which was characterized by altered DH-VH and reduced VH-PL, VH-BLA, and PL-BLA synchrony. This reduction in VH-PL-BLA synchrony was not fully accounted for by non-specific locomotor effects, as revealed by comparing between epochs of movement and freezing in the controls. These results suggest that D1Rs regulate CFC by modulating synchronized oscillations within the hippocampus-prefrontal-amygdala circuit. They also add to growing evidence indicating that this circuit synchrony at retrieval reflects a neural signature of learned fear.

## Introduction

Learning that certain environments predict threat is adaptive and can be investigated using contextual fear conditioning (CFC) in rodents. During CFC, unsignalled presentation of an aversive unconditioned stimulus (US; e.g. footshock) typically occurs in a novel context. This results in the encoding of a representation of the context, which becomes associated with the US. Fear-related behavior (e.g. freezing) is then elicited in the conditioned context during later memory retrieval [Rudy et al., 2004]. CFC provides a useful model for studying the neurobiological mechanisms underpinning emotional learning, which is also translationally relevant since aberrant emotional memory processing is a key feature of various anxiety-related disorders [Maren et al., 2013; Chaaya et al., 2018].

CFC requires coordinated activity within a distributed neural network that includes the hippocampus, amygdala, and prefrontal cortex (PFC). The context representation is thought to be encoded in dorsal hippocampus (DH) and conveyed to basolateral amygdala (BLA) for association with the US representation [Rudy et al., 2004; Fanselow, 2010; Maren et al., 2013; Ohkawa et al., 2015]. Other evidence indicates that DH is also involved in forming and storing the context-US association [Chang et al. 2008; Chaaya et al., 2018]. Prelimbic (PL) PFC plays a role in processing the context and US representations through its inter-connections with DH and BLA [Zelikowsky et al., 2014; Kitamura et al., 2017; Shibano et al., 2020]. Importantly, DH projects to BLA and PL indirectly through ventral hippocampus (VH), which is also involved in encoding representations of the context and US via its inter-connections with BLA and PL [Pitkänen et al., 2000; Cenquizca & Swanson 2007; Huff et al., 2016; Kim & Cho, 2017, 2020; Jimenez et al., 2020; Twining et al., 2020].

In contrast to the neural circuit basis of CFC, its regulation by neuromodulators remains unclear [Likhtik & Johansen, 2019]. Dopamine regulates CFC via D1-like receptor (D1R) signalling in the hippocampal-prefrontal-amygdala circuit [Stubbendorff & Stevenson, 2021]. These brain areas receive midbrain dopamine projections and express D1Rs [Oades & Halliday, 1987; Gasbarri et al., 1997; Missale et al., 1998]. Systemic D1R blockade impairs the acquisition of contextual fear [Inoue et al., 2000; Heath et al., 2015; Stubbendorff et al., 2019a]. In terms of the brain areas involved, blocking D1Rs locally in DH, PL, or BLA, but not VH, disrupts CFC [Heath et al., 2015; Stubbendorff et al., 2019a]. D1R blockade also interferes with long-term potentiation (LTP), a model of synaptic plasticity, within this circuitry [Huang & Kandel, 1995; Gurden et al. 2000; Li et al., 2011]. However, the neurophysiological mechanisms linking D1R modulation of LTP and CFC remain poorly understood.

Synaptic plasticity underpinning learning is facilitated by synchronized rhythmic oscillations that mediate communication between inter-connected brain areas. Theta and gamma synchrony and cross-frequency coupling in the hippocampal-prefrontal-amygdala circuit are important for learned fear processing [Headley & Paré, 2013; Stujenske et al., 2014; Bocchio et al., 2017; Makino et al., 2019; Radiske et al., 2020; Totty & Maren, 2022]. Interestingly, D1Rs modulate oscillatory activity in and synchrony between these areas [Bo & Savoldi, 1992; Weiss et al., 2003; Lorétan et al., 2004; Parker et al., 2014; Xu et al., 2016; Ott et al., 2018; Park et al., 2021; Gonzalez et al., 2021], raising the possibility that D1R blockade impairs LTP and CFC by disrupting synchronized oscillations in this circuit.

In this study we examined the effects of systemic D1R blockade on theta and gamma oscillations in DH, VH, PL, and BLA during CFC. Since synchronized oscillations and cross-frequency coupling in this circuit play a role in fear memory retrieval [Headley & Paré, 2013; Stujenske et al., 2014; Bocchio et al., 2017; Makino et al., 2019; Totty & Maren, 2022], we also examined the effects of impaired CFC by D1R blockade on oscillatory synchrony between these areas at retrieval. We found that blocking D1Rs altered DH-VH and reduced VH-PL and VH-BLA synchrony acutely during conditioning. We also found that impaired CFC by D1R blockade was associated with altered DH-VH and reduced VH-PL-BLA synchrony during later retrieval testing.

## Results

### D1R blockade impairs CFC

To determine if D1Rs regulate CFC, we examined the effects of the selective D1R antagonist SCH23390 given systemically (0.1 mg/kg, i.p.) before CFC on freezing at retrieval (Fig 1A-1C). SCH23390 (n=20) impaired CFC, as indicated by reduced freezing during retrieval testing, compared to vehicle-treated controls (n=22). For freezing throughout the test session (Fig 1B), an unpaired t-test revealed that SCH23390 resulted in decreased freezing, compared to vehicle (t(40)=2.65, P=0.012). This was confirmed by the time course analysis (Fig 1C), which showed that SCH23390 resulted in decreased freezing throughout retrieval, compared to vehicle. Two-way ANOVA revealed a significant main effect of treatment (F(1,40)=7.01, P=0.012) but no main effect of time (F(4,160)=2.32, P=0.059) or treatment x time interaction (F(4,160)=0.74, P=0.57). This confirms previous results showing that SCH23390 impairs CFC [Inoue et al., 2000; Heath et al., 2015; Stubbendorff et al., 2019a].

**Fig 1.**
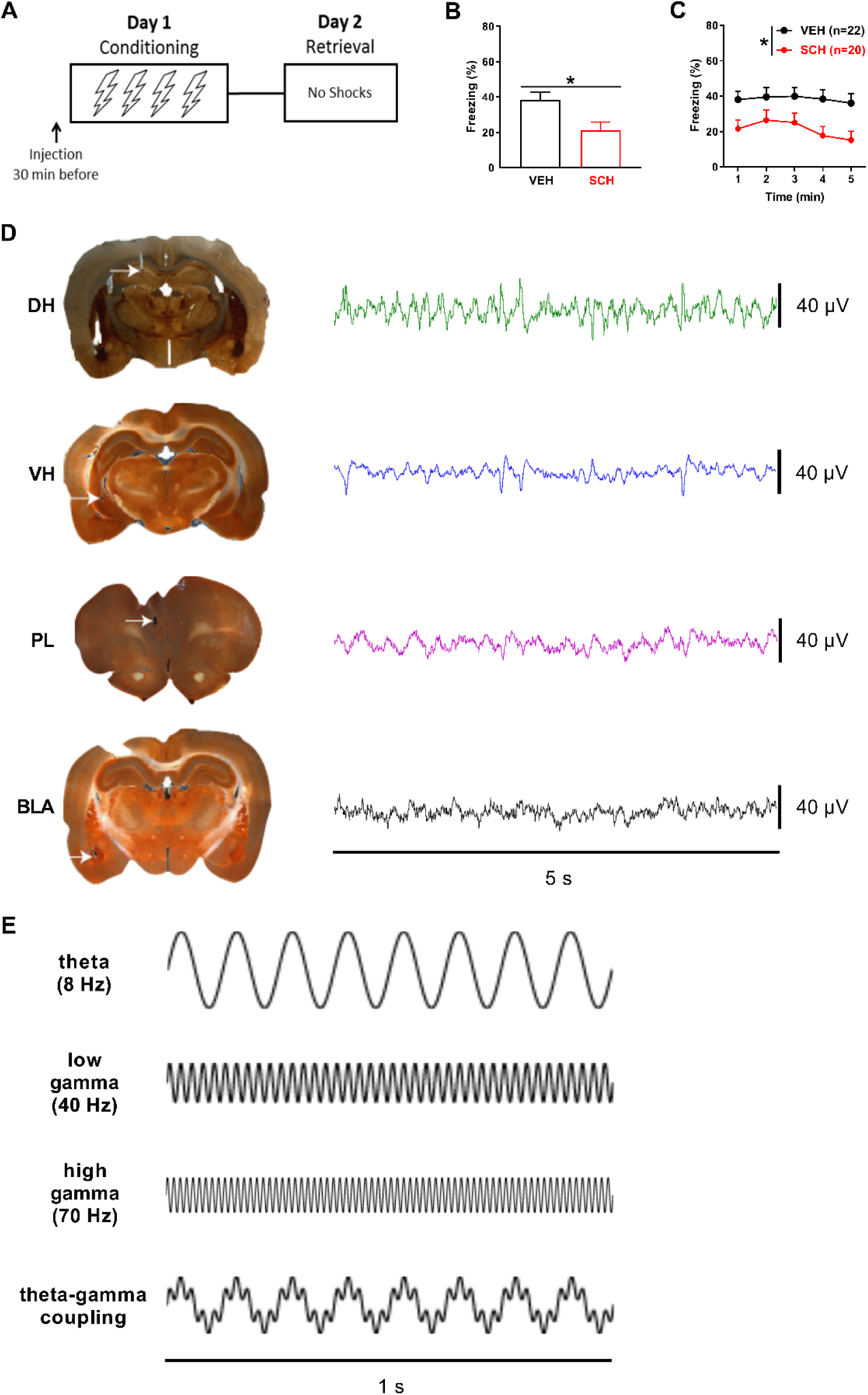
D1R blockade impairs CFC. A) Schematic representation of the behavioral testing paradigm used. Rats received a systemic injection of SCH23390 (SCH) or vehicle (VEH) 30 min before conditioning, and LFPs were recorded during conditioning and retrieval testing. B) SCH given before CFC decreased freezing at retrieval, compared to VEH (**\***P<0.05). C) A time course analysis showed that SCH resulted in decreased freezing throughout retrieval, compared to VEH (**\***P<0.05). D) Illustrative examples of electrode placements in (left; white arrows) and LFP signals recorded from (right) DH, VH, PL, and BLA. E) Schematic representations of theta (8 Hz), low gamma (40 Hz), and high gamma (70 Hz) oscillations, and theta-gamma cross-frequency coupling.

Examples of electrode placements in and local field potentials (LFPs) recorded from each area before US presentations during CFC are shown in Fig 1D. Schematic representations of theta (4-12 Hz), low gamma (30-45 Hz), and high gamma (55-80 Hz) oscillations, and theta-gamma coupling, are shown in Fig 1E. Compared to the behavioral data analysis, fewer rats were included in the electrophysiological data analyses because of missed electrode placements or electrical noise artefacts contained in the data. For the analyses comparing between epochs of movement and freezing at retrieval in vehicle-treated controls, the numbers of controls included in these comparisons were unequal because some animals had no movement or freezing epochs based on the criteria defined for their inclusion in the analyses (see Methods). The numbers of animals included in the final datasets for each comparison (SCH23390 vs vehicle treatment; movement vs freezing in vehicle-treated controls) are indicated in Figs 2-7.

**Fig 2.**
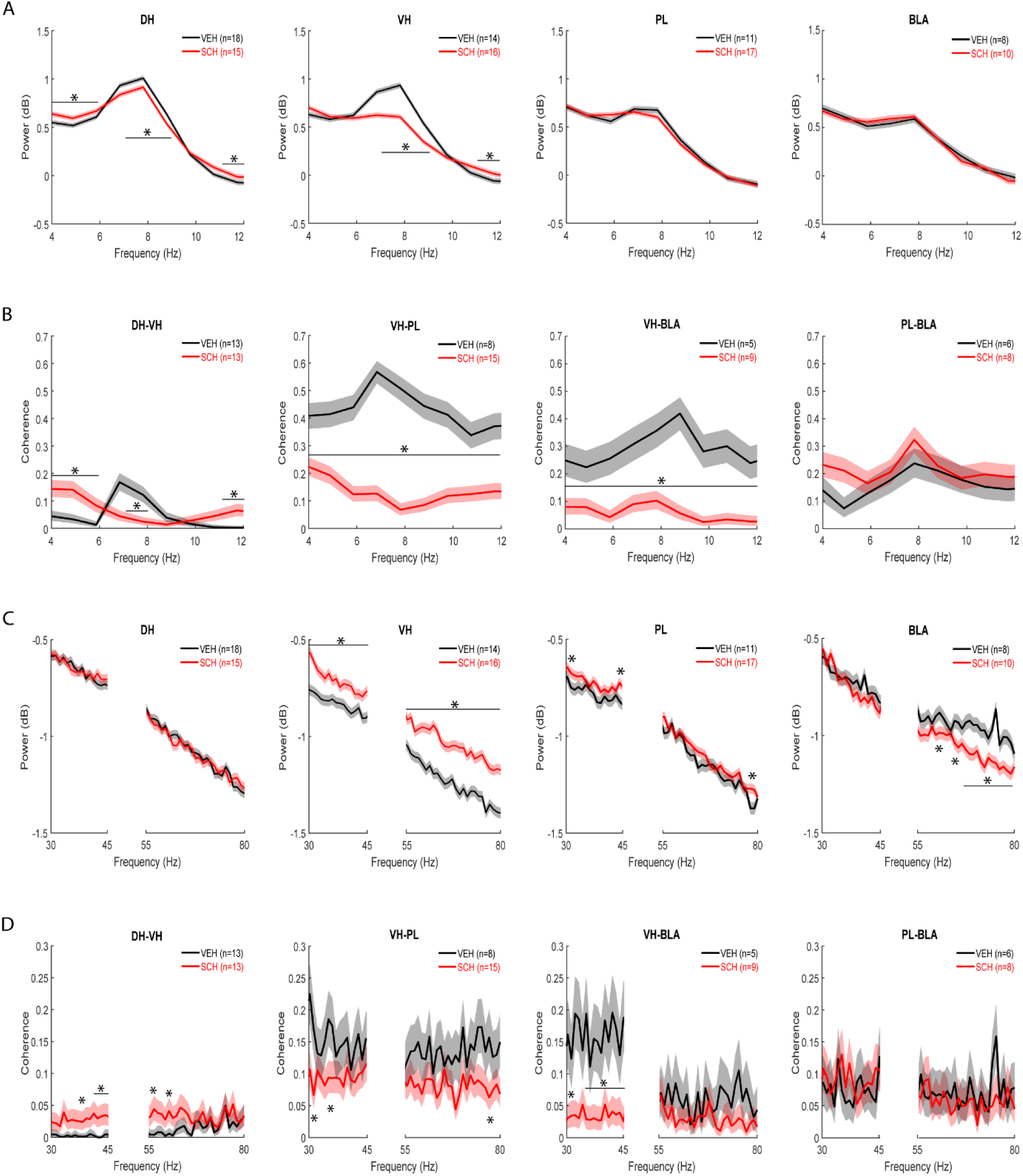
D1R blockade alters DH-VH and reduces VH-PL and VH-BLA synchrony during CFC. A) Acute effects of SCH23390 (SCH) on theta power during CFC. Peak theta power occurred ∼7-8 Hz in controls treated with vehicle (VEH). Compared to VEH, SCH decreased peak theta power and increased power outside the peak in DH and VH (*P<0.001), but not PL or BLA. B) Acute effects of SCH on theta coherence during CFC. Peak theta coherence occurred ∼7-9 Hz with VEH. Compared to VEH, SCH decreased DH-VH peak theta coherence and increased DH-VH coherence outside the peak (*P<0.001). SCH decreased VH-PL and VH-BLA theta coherence, compared to VEH (*P<0.001), without affecting PL-BLA theta coherence. C) Acute effects of SCH on gamma power during CFC. SCH had no effect on gamma power in DH. Compared to VEH, SCH increased gamma power in VH and PL (*P<0.001). SCH decreased high gamma power in BLA, compared to VEH (*P<0.001). D) Acute effects of SCH on gamma coherence before CFC. SCH increased DH-VH and decreased VH-PL gamma coherence, compared to VEH (*P<0.001). SCH decreased VH-BLA low gamma coherence, compared to VEH (*P<0.001), without affecting PL-BLA gamma coherence.

### Acute D1R blockade alters intra-hippocampal and reduces hippocampal-prefrontal and hippocampal-amygdala synchrony during conditioning

To determine if D1Rs modulate theta oscillations in the hippocampus-prefrontal-amygdala circuit during CFC, we examined the acute effects of SCH23390 on theta power in each area in the 2 min period before US presentations during conditioning (Fig 2A). Differences between SCH23390 and vehicle treatment in power at each individual frequency throughout the theta band were quantified using a log ratio test. In vehicle-treated controls there was a prominent peak ∼7-8 Hz in hippocampus, whereas this peak was less pronounced in PL and BLA. In DH, SCH23390 decreased peak theta power (7-9 Hz) and increased power outside the peak (4-6 and 11-12 Hz), compared to vehicle (P<0.001). The largest effect of SCH23390 on theta power was observed in VH, where it reduced peak power (7-9 Hz) while also increasing power at higher frequencies (11-12 Hz), compared to vehicle (P<0.001). SCH23390 had no effect on theta power in PL or BLA.

To determine if D1Rs modulate theta synchrony during CFC, we examined the acute effects of SCH23390 on theta coherence in this circuit (Fig 2B). Differences between SCH23390 and vehicle treatment in coherence between the areas sharing direct anatomical connections (i.e. DH-VH, VH-PL, VH-BLA, and PL-BLA) were quantified at each individual frequency throughout the theta band using a chi-squared difference of coherence test. Vehicle-treated controls showed peaks for theta coherence ∼7-9 Hz. SCH23390 decreased peak theta coherence between DH and VH (7-8 Hz), while increasing coherence outside the peak (4-6 and 11-12 Hz), compared to vehicle (P<0.001). The largest effects of SCH23390 were observed on VH-PL and VH-BLA theta coherence, where SCH23390 decreased coherence throughout the theta band, compared to vehicle (P<0.001). SCH23390 had no effect on theta coherence between PL and BLA.

To determine if D1Rs modulate gamma oscillations in this circuit during CFC, we examined the acute effects of SCH23390 on gamma power in each area (Fig 2C). Differences between SCH23390 and vehicle treatment in power at each individual frequency throughout the low and high gamma bands were quantified using a log ratio test. No obvious peak frequencies were observed for gamma power in any area in vehicle-treated controls. In DH, there was no effect of SCH23390 on gamma power. The largest effect of SCH23390 on gamma power was observed in VH, where power was increased throughout the low and high gamma bands, compared to vehicle (P<0.001). In PL, SCH23390 increased power at some low (31-32 and 44-45 Hz) and high (78-79 Hz) gamma frequencies, compared to vehicle (P<0.001). In BLA, SCH23390 had no effect on low gamma power and decreased high gamma power at various frequencies (60-61, 64-65, and 67-79 Hz, compared to vehicle (P<0.001).

To determine if D1Rs modulate gamma synchrony during CFC, we examined the acute effects of SCH23390 on gamma coherence in this circuit (Fig 2D). Differences between SCH23390 and vehicle treatment in coherence were quantified at each individual frequency throughout the low and high gamma bands using a chi-squared difference of coherence test. There were no obvious peak frequencies observed for gamma coherence in vehicle-treated controls. SCH23390 increased coherence between DH and VH at low (37-39 and 41-45 Hz) and high (56-57 and 60-62 Hz) gamma frequencies, compared to the low levels of gamma coherence observed with vehicle (P<0.001). In contrast, coherence between VH and PL was decreased by SCH23390 at some low (30-31 and 35-36 Hz) and high (74-75 Hz) gamma frequencies, compared to vehicle (P<0.001). SCH23390 also decreased gamma coherence between VH and BLA at low (30-33 and 35-45 Hz; P<0.001) but not high gamma frequencies, compared to vehicle. SCH23390 had no effect on gamma coherence between PL and BLA, compared to vehicle.

To determine if D1Rs modulate theta-gamma coupling in this circuit during CFC, we examined the acute effects of SCH23390 on theta-gamma phase-amplitude coupling (PAC) in each area (Fig 3A- B). Theta-gamma PAC was quantified using a modulation index. Although peak theta power in each area occurred ∼7-8 Hz (Fig 2A), color plots of theta-gamma PAC in each area showed that peak theta phase coupling of gamma amplitude occurred ∼5 Hz with vehicle (Fig 3A) or SCH23390 (not shown). Therefore we focused the analysis on this theta frequency. The quantitative analysis using two-way ANOVA found no acute effects of SCH23390 on theta-gamma PAC in any area (Fig 3B and Table S1).

**Fig 3.**
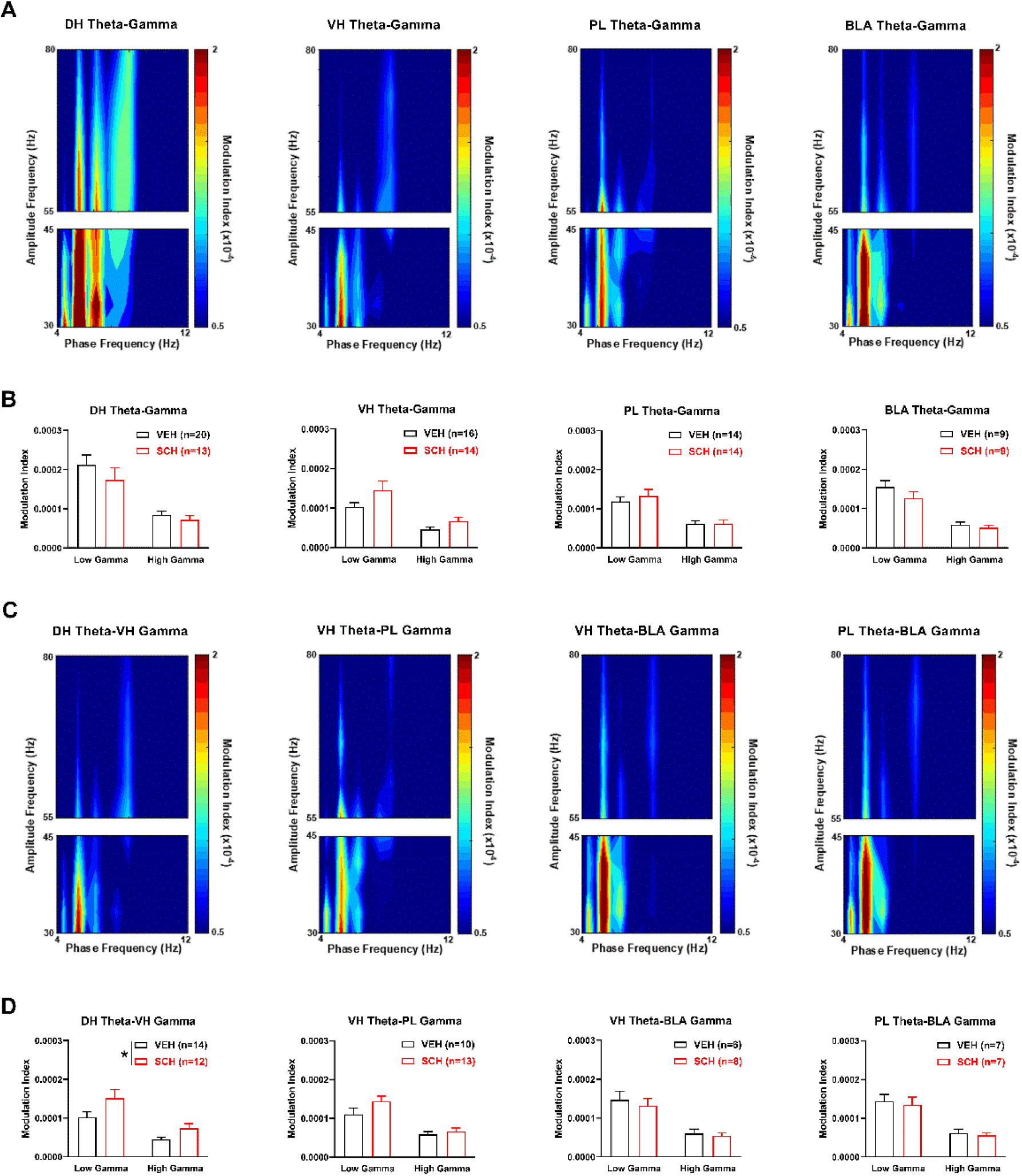
D1R blockade enhances theta-gamma coupling between DH and VH during CFC. A) Theta-gamma PAC in each area after vehicle (VEH) treatment, with blue and red indicating lower and higher gamma amplitude, respectively. Peak PAC occurred at a theta frequency ∼5 Hz in each area. B) SCH23390 (SCH) had no effects on theta-gamma PAC in DH, VH, PL or BLA. C) Theta-gamma PAC between areas with VEH, showing that peak PAC occurred ∼5 Hz. D) SCH increased DH theta coupling of VH gamma, compared to VEH (**\***P<0.05). SCH had no effects on VH theta-PL gamma, VH theta-BLA gamma, or PL theta-BLA gamma PAC.

We also examined the acute effects of SCH23390 on theta-gamma PAC between the directly inter-connected areas during CFC (Fig 3C-D). Color plots of theta-gamma PAC between areas showed that peak PAC occurred ∼5 Hz with vehicle (Fig 3C) or SCH23390 (not shown), therefore the analysis was focused on this theta frequency. Compared to vehicle, SCH23390 increased DH theta coupling of VH gamma (Fig 3D). Two-way ANOVA revealed a significant main effect of treatment (F(1,24)=4.41, P=0.046) but no treatment x frequency interaction (F(1,24)=1.75, P=0.20). For VH theta coupling of PL gamma, two-way ANOVA found no main effect of treatment (F(1,21)=1.61, P=0.22) but revealed a significant treatment x frequency interaction (F(1,21)=4.53, P=0.045); however, post-hoc testing found no difference between vehicle and SCH23390 treatment at low or high gamma frequencies. There were also no effects of SCH23390 on VH or PL theta coupling of BLA gamma (Table S1).

### Impaired CFC by D1R blockade is associated with altered intra-hippocampal and reduced hippocampal-prefrontal-amygdala synchrony at retrieval

To determine a role for hippocampal-prefrontal-amygdala theta oscillations in contextual fear memory, we examined the effects of impaired CFC by SCH23390 on theta power in each area during later retrieval tested drug-free (Fig 4A). Compared to theta power during CFC, there was less of a peak that was shifted to 6-7 Hz in hippocampus, with much less prominent peaks in PL or BLA, in vehicle-treated controls. Overall, SCH23390 shifted the peak to 7-8 Hz in each area. SCH23390 resulted in the biggest increase in peak theta power in DH (7-8 Hz), while also decreasing power at lower frequencies (4-6 Hz), compared to vehicle (P<0.001). In VH, SCH23390 decreased power at lower (5-7 Hz) and increased power at higher (11-12 Hz) theta frequencies, compared to vehicle (P<0.001). SCH23390 increased peak theta power in PL (8-9 Hz) and BLA (7-9 Hz), compared to vehicle (P<0.001).

**Fig 4.**
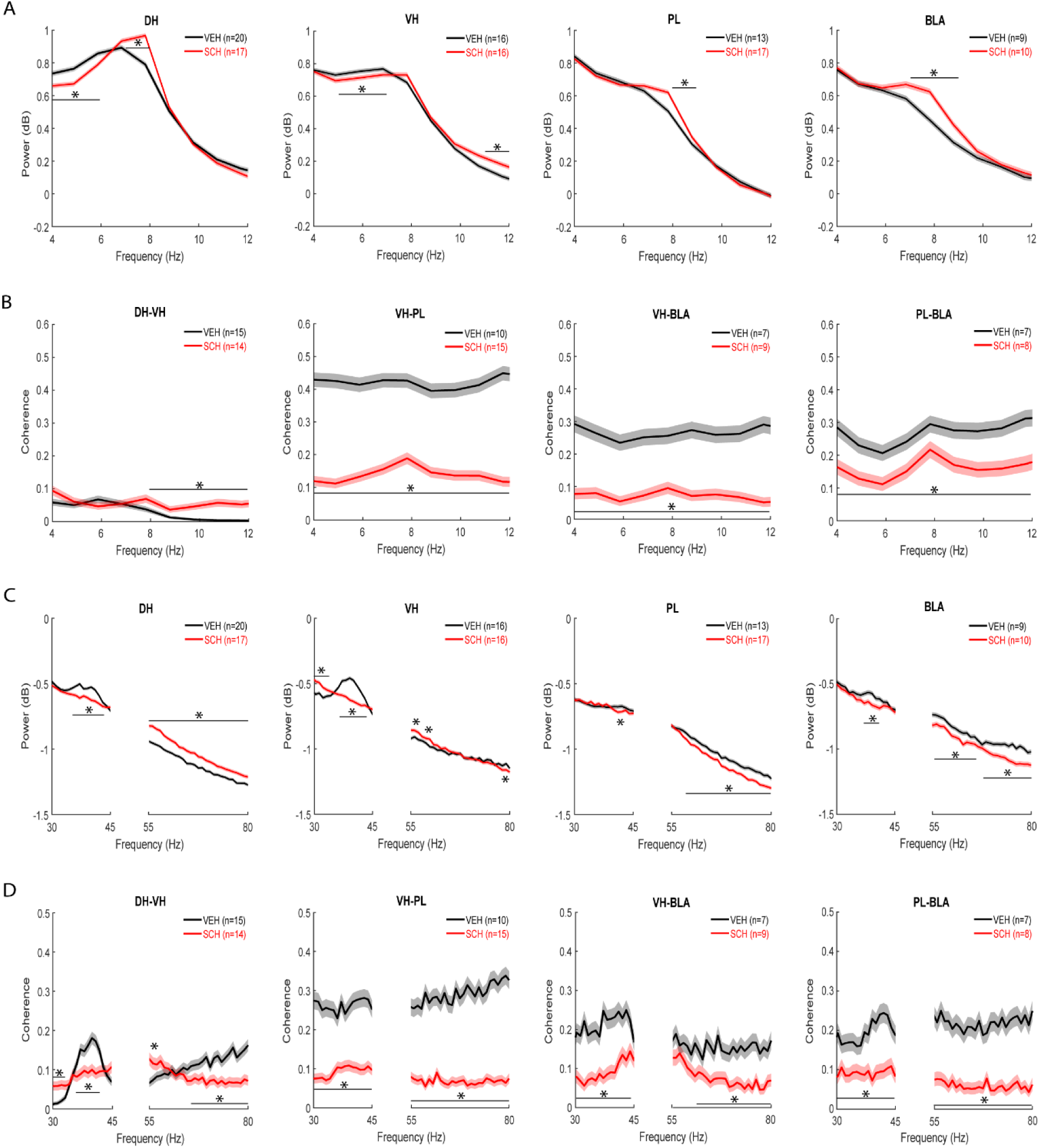
Impaired CFC by D1R blockade is associated with altered DH-VH and reduced VH-PL-BLA synchrony at retrieval. A) Effects of SCH23390 (SCH) given before CFC on theta power at retrieval. Peak theta power occurred ∼6-7 Hz in controls treated with vehicle (VEH), while SCH shifted the peak to ∼7-8 Hz. In DH, SCH decreased lower and increased peak theta power, compared to VEH (*P<0.001). In VH, SCH decreased lower and increased higher theta power, compared to VEH (*P<0.001). In PL and BLA, SCH increased theta power, compared to VEH (*P<0.001). B) Effects of SCH given before CFC on theta coherence at retrieval. SCH increased DH-VH and decreased VH-PL, VH-BLA, and PL-BLA theta coherence, compared to VEH (*P<0.001). C) Effects of SCH given before CFC on gamma power at retrieval. A peak in low gamma power occurred ∼35-40 Hz in DH, VH, and BLA, but not PL, with VEH. In DH, SCH decreased low and increased high gamma power, compared to VEH (*P<0.001). In VH, SCH decreased peak low gamma power and increased low gamma power outside the peak, while increasing or decreasing power at various high gamma frequencies, compared to VEH (*P<0.001). In PL and BLA, SCH decreased gamma power, compared to VEH (*P<0.001). D) Effects of SCH given before CFC on gamma coherence at retrieval. Peak low gamma coherence between DH and VH occurred at 35-40 Hz with VEH, but no peaks were observed for coherence between the other areas. SCH decreased DH-VH peak low gamma coherence and increased low gamma coherence outside the peak, while increasing or decreasing coherence at various high gamma frequencies, compared to VEH (*P<0.001). SCH decreased VH-PL, VH-BLA, and PL-BLA gamma coherence, compared to VEH (*P<0.001).

To determine a role for theta synchrony in contextual fear memory, we examined the effects of impaired CFC by SCH23390 on theta coherence in this circuit at retrieval (Fig 4B). Compared to theta power, peaks for theta coherence were less pronounced or absent. Overall, SCH23390 increased theta coherence within hippocampus and decreased theta coherence between VH, PL, and BLA. SCH23390 increased coherence between DH and VH at higher theta frequencies (8-12 Hz), compared to vehicle (P<0.001). In contrast, SCH23390 caused a large decrease in VH-PL, VH-BLA, and PL-BLA coherence throughout the theta band, compared to vehicle (P<0.001).

To determine a role for gamma oscillations in contextual fear memory, we examined the effects of impaired CFC by SCH23390 on gamma power in each area at retrieval (Fig 4C). Compared to low gamma power during CFC, there was a peak at 35-40 Hz in hippocampus and, to a lesser extent, in BLA and PL in vehicle-treated controls. In DH, SCH23390 decreased peak low gamma power (35-43 Hz) and increased power throughout the high gamma band, compared to vehicle (P<0.001). In VH, SCH23390 decreased peak low gamma power (36-43 Hz) but increased low gamma power outside the peak (30-34 Hz), while increasing power at some (55-57 and 59-60 Hz) and decreasing power at other (78-79 Hz) high gamma frequencies, compared to vehicle (P<0.001). In PL, SCH23390 decreased power at some low (41-42 Hz) and most high (58-80 Hz) gamma frequencies, compared to vehicle (P<0.001). In BLA, SCH23390 decreased peak low gamma power (37-41 Hz) and high gamma power at most frequencies (55-66 and 68-80 Hz), compared to vehicle (P<0.001).

To determine a role for gamma synchrony in contextual fear memory, we examined the effects of impaired CFC by SCH23390 on gamma coherence in this circuit at retrieval (Fig 4D). Vehicle-treated controls showed a peak for low gamma coherence within hippocampus at 35-40 Hz but there were no obvious peaks observed for gamma coherence between any other areas. SCH23390 decreased peak low gamma coherence between DH and VH (36-42 Hz) and increased low gamma coherence outside the peak (30-33 Hz), while increasing coherence at some (55-56 Hz) and decreasing coherence at other (65- 80 Hz) high gamma frequencies, compared to vehicle (P<0.001). SCH23390 caused a large decrease in coherence between VH and PL throughout the low and high gamma bands, compared to vehicle (P<0.001). SCH23390 also decreased coherence between VH and BLA at most low (30-44 Hz) and high (61-80 Hz) gamma frequencies, compared to vehicle (P<0.001). Similarly, SCH23390 resulted in a large decrease in coherence between PL and BLA across the low and high gamma bands, compared to vehicle (P<0.001).

To determine a role for theta-gamma coupling in contextual fear memory, we examined the effects of impaired CFC by SCH23390 on theta-gamma PAC in each area at retrieval (Fig 5A-B). Color plots of theta-gamma PAC in each area showed that peak PAC occurred ∼5 Hz with vehicle (Fig 5A) or SCH23390 (not shown), thus we focused the analysis on this frequency. The quantitative analysis found no effects of impaired CFC by SCH23390 on theta-gamma PAC in any area at retrieval (Fig 5B and Table S2).

**Fig 5.**
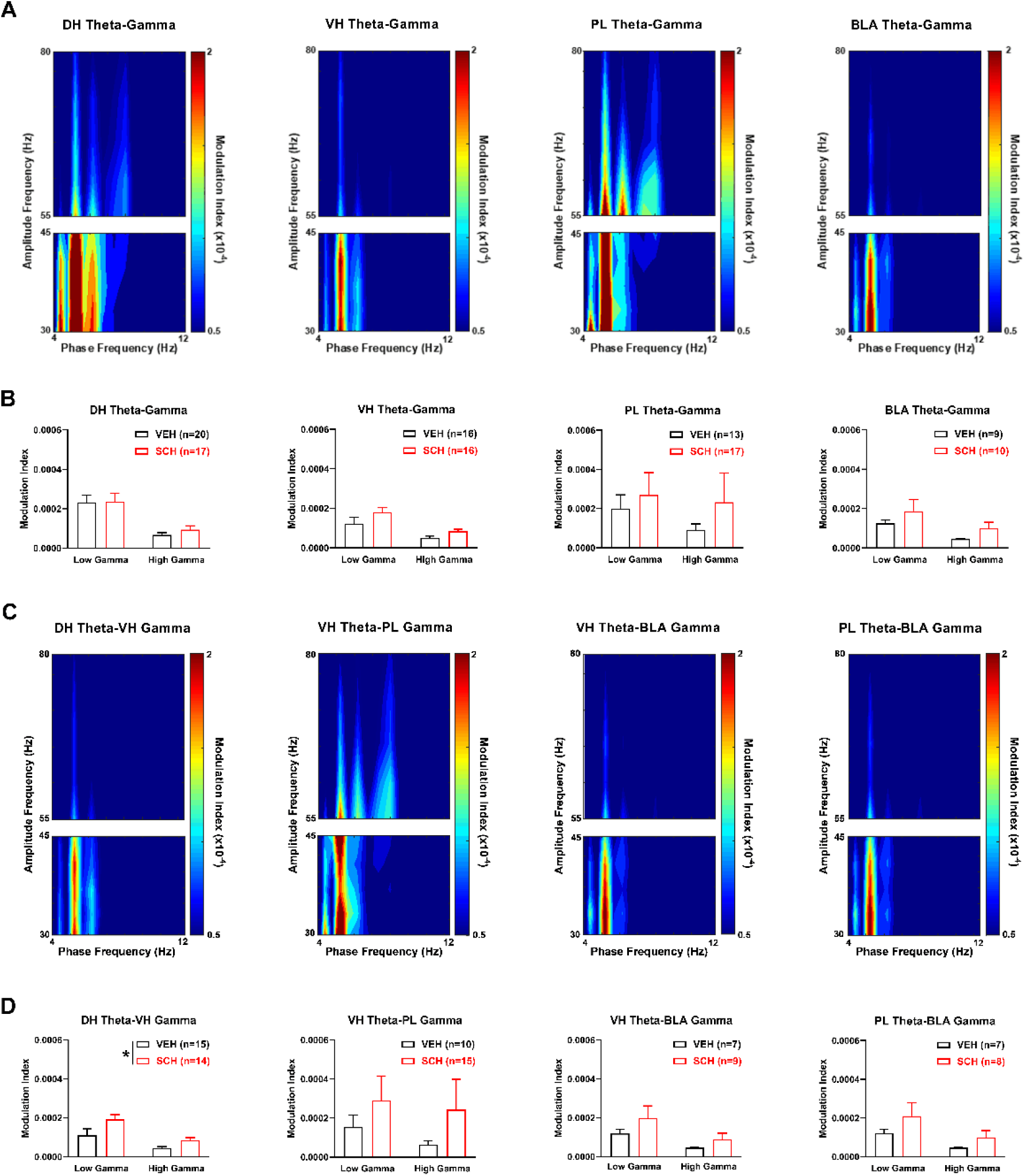
Impaired CFC by D1R blockade is associated with enhanced theta-gamma coupling between DH and VH at retrieval. A) Theta-gamma PAC in each area after vehicle (VEH) treatment, with lower and higher gamma amplitude indicated by blue and red, respectively. Peak PAC occurred ∼5 Hz in each area. B) SCH23390 (SCH) had no effects on theta-gamma PAC in DH, VH, PL or BLA. C) Theta-gamma PAC between areas with VEH, showing that peak PAC occurred ∼5 Hz. D) SCH increased DH theta phase coupling of gamma amplitude in VH, compared to VEH (**\***P<0.05). SCH had no effects on VH theta-PL gamma, VH theta-BLA gamma, or PL theta- BLA gamma PAC.

We also examined the effects of impaired CFC by SCH2330 on theta-gamma PAC between these areas at retrieval (Fig 5C-D). Color plots of theta-gamma PAC between areas showed that peak PAC occurred ∼5 Hz with vehicle (Fig 5C) or SCH23390 (not shown), therefore we focused the analysis on this frequency. Compared to vehicle, SCH23390 increased DH theta coupling of VH gamma (Fig 5D). Two-way ANOVA revealed a significant main effect of treatment (F(1,27)=5.03, P=0.033) but no treatment x frequency interaction (F(1,27)=1.70, P=0.20). SCH23390 had no effects on VH theta coupling of gamma in PL or BLA; similarly, there was no effect of SCH23390 on PL theta coupling of BLA gamma (Table S2).

### Reduced hippocampal-prefrontal-amygdala synchrony associated with impaired CFC by D1R blockade is not fully accounted for by increased movement at retrieval

Because theta and gamma oscillations in these areas are associated with movement [Amir et al., 2018; Foo & Bohbot, 2020; Nuñez & Buño, 2021], and contextual fear was inferred from freezing, we also compared synchronized oscillations between epochs of movement and freezing during retrieval testing in the vehicle-treated controls. We reasoned that if the reduced freezing and associated oscillatory synchrony resulting from SCH23390 given before CFC simply reflected increased movement at retrieval then we would find a similar synchrony profile with movement, compared to freezing, in the controls at retrieval. However, if we found differences between these synchrony profiles then this may instead reflect different neural signatures for movement and contextual fear memory.

In terms of hippocampal-prefrontal-amygdala theta oscillations in the controls at retrieval, movement was associated with a peak in theta power at 7-8 Hz in each area (Fig 6A). In DH, movement increased peak theta power (7-9 Hz) and decreased theta power outside the peak (4-6 and 11-12 Hz), compared to freezing (P<0.001). In VH, movement also increased peak theta power (8-9 Hz), while theta power was also decreased outside the peak (4-6 and 11-12 Hz), compared to freezing (P<0.001). Movement increased peak theta power in PL (8-9 Hz) and BLA (7-9 Hz), compared to freezing (P<0.001).

**Fig 6.**
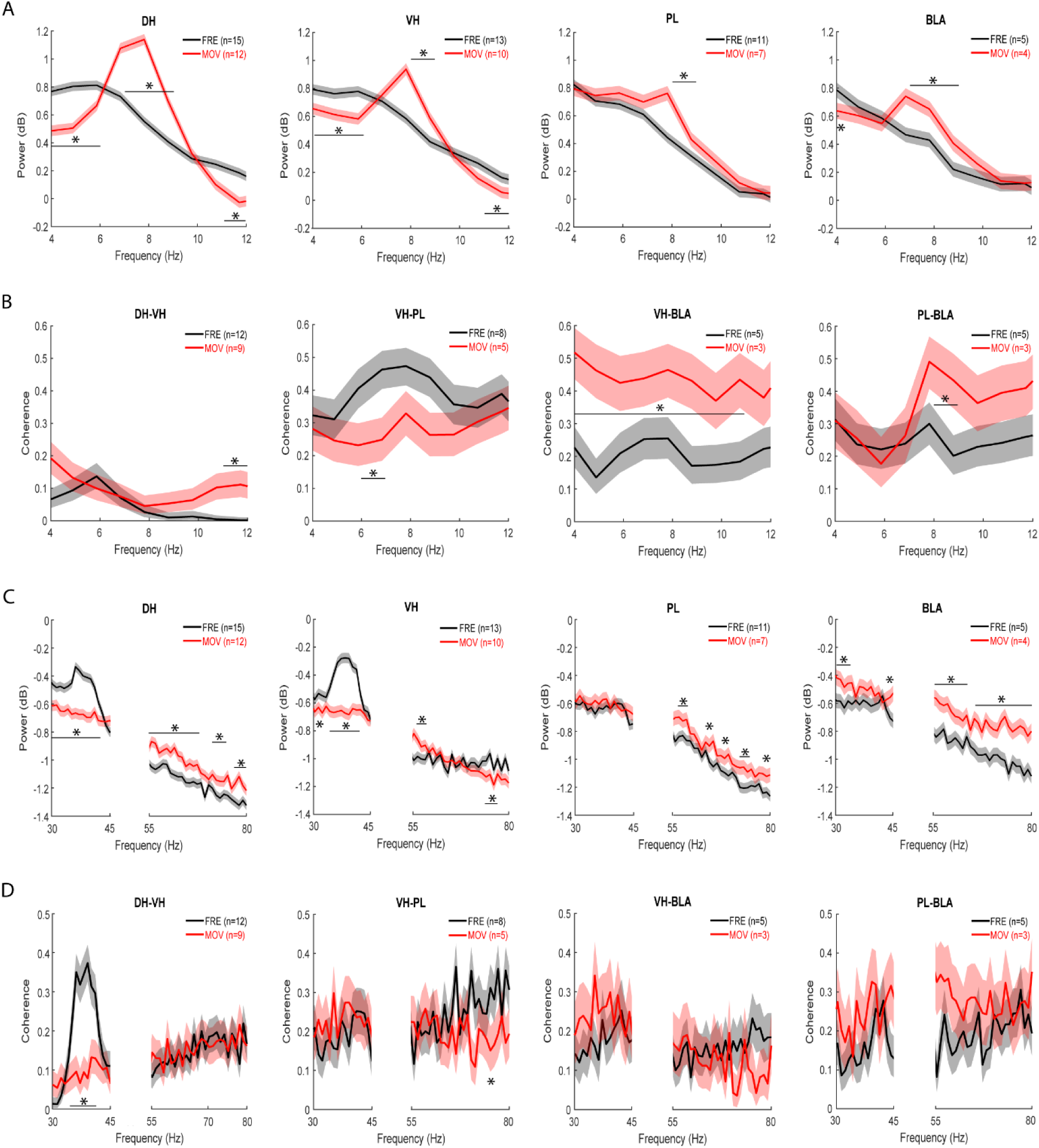
Movement and freezing in vehicle-treated controls are associated with differences in DH-VH and VH-PL- BLA synchrony at retrieval. A) Differences between movement (MOV) and freezing (FRE) in theta power at retrieval. Peak theta power occurred ∼7-8 Hz with MOV. Compared to FRE, MOV increased peak theta power and decreased theta power outside the peak in DH and VH (*P<0.001). In PL and BLA, MOV increased theta power, compared to FRE (*P<0.001). B) Differences between MOV and FRE in theta coherence at retrieval. Compared to FRE, MOV increased DH-VH, VH-BLA, and PL-BLA theta coherence (*P<0.001). MOV decreased VH-PL theta coherence, compared to FRE (*P<0.001). C) Differences between MOV and FRE in gamma power at retrieval. A peak in low gamma power occurred ∼35-40 Hz with FRE in DH and VH, but not PL or BLA. In DH, MOV decreased low and increased high gamma power, compared to FRE (*P<0.001). In VH, MOV decreased low gamma power, while power was increased or decreased at various high gamma frequencies, compared to FRE (*P<0.001). In PL, MOV increased high gamma power, compared to FRE (*P<0.001). In BLA, MOV increased gamma power, compared to FRE (*P<0.001). D) Differences between MOV and FRE in gamma coherence at retrieval. Peak low gamma coherence between DH and VH occurred ∼35-40 Hz with FRE, but there were no peaks for gamma coherence between other areas. MOV decreased DH-VH low gamma coherence, compared to FRE (*P<0.001). MOV decreased VH-PL high gamma coherence, compared to FRE (*P<0.001). MOV had no effects on VH-BLA or PL-BLA gamma coherence.

In terms of theta synchrony in this circuit in the controls at retrieval, there were less obvious peaks in theta coherence with movement in comparison to theta power (Fig 6B). Movement increased coherence between DH and VH at higher theta frequencies (11-12 Hz), compared to freezing (P<0.001). In contrast, movement decreased peak theta coherence (6-7 Hz) between VH and PL, compared to freezing (P<0.001). Movement increased coherence between VH and BLA across most theta frequencies (4-11 Hz), compared to freezing (P<0.001). Movement also increased peak theta coherence (8-9 Hz) between PL and BLA, compared to freezing (P<0.001).

In terms of gamma oscillations in this circuit in the controls at retrieval, freezing was associated with a peak in low gamma power at 35-40 Hz in hippocampus, but not in PL or BLA (Fig 6C). In DH, movement decreased low (30-42 Hz) and increased high (55-67, 71-75, and 77-80 Hz) gamma power, compared to freezing (P<0.001). In VH, movement also decreased low gamma power (30-32 and 34- 42 Hz), while power was increased (56-59 Hz) or decreased (74-77 Hz) at various high gamma frequencies, compared to freezing (P<0.001). In PL, movement had no effect on low and increased high gamma power (56-59, 64-65, 67-69, 72-75, and 78-80), compared to freezing (P<0.001). In BLA, movement increased both low (30-34 and 44-45 Hz) and high (55-63 and 65-80 Hz) gamma power, compared to freezing (P<0.001).

In terms of gamma synchrony in this circuit in the controls at retrieval, freezing was associated with a peak in low gamma coherence at 35-40 Hz within hippocampus but not between any other areas (Fig 6D). Movement abolished peak low gamma coherence between DH and VH (35-41 Hz; P<0.001) but had no effect on high gamma coherence, compared to freezing. Movement had no effect on low gamma coherence between VH and PL but decreased coherence at some high gamma frequencies (70- 71 Hz), compared to freezing (P<0.001). Movement had no effect on gamma coherence between VH and BLA, or between PL and BLA.

In terms of theta-gamma coupling in this circuit in the controls at retrieval, color plots of theta-gamma PAC in each area showed that peak PAC occurred ∼5 Hz with freezing (Fig 7A) or movement (not shown), therefore the analysis was focused on this frequency. The quantitative analysis found no differences between movement and freezing in DH, VH, or PL (Fig 7B and Table S3). However, compared to freezing, movement increased theta coupling of low gamma in BLA. Two-way ANOVA showed no main effect of locomotion (F(1,7)=4.05, P=0.084) but revealed a marginally significant locomotion x frequency interaction (F(1,7)=5.59, P=0.050). Post-hoc testing confirmed that theta coupling of low gamma was significantly increased with movement, compared to freezing (P<0.05).

**Fig 7.**
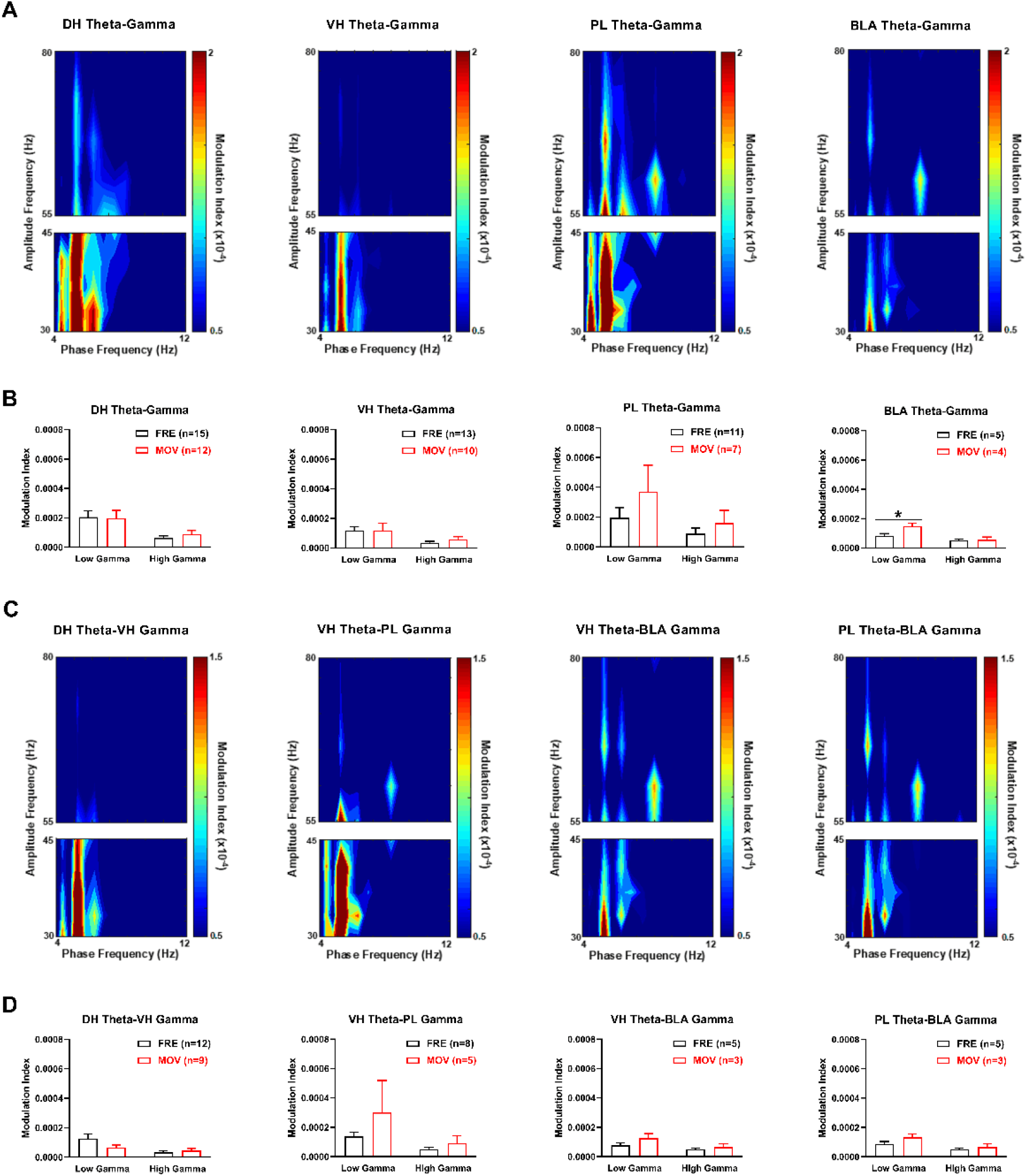
Movement in vehicle-treated controls enhances theta-gamma coupling in BLA at retrieval. A) Theta-gamma PAC in each area during freezing (FRE), with lower and higher gamma amplitude indicated by blue and red, respectively. Peak PAC occurred ∼5 Hz in each area. B) Movement (MOV) had no effect on theta-gamma PAC in DH, VH, or PL. MOV increased theta coupling of low gamma in BLA, compared to FRE (*P<0.05). C) Theta- gamma PAC between areas during FRE, showing that peak PAC occurred ∼5 Hz. D) MOV had no effects on DH theta-VH gamma, VH theta-PL gamma, VH theta-BLA gamma, or PL theta-BLA gamma PAC.

We also examined differences between movement and freezing in theta-gamma coupling between these areas in the controls at retrieval (Fig 7C-D). Color plots of theta-gamma PAC between areas showed that peak PAC occurred ∼5 Hz with freezing (Fig 7C) or movement (not shown), therefore we focused our analysis at this frequency. The quantitative analysis found no differences between movement and freezing on theta-gamma PAC between any areas at retrieval (Fig 7D and Table S3). For DH theta coupling of VH gamma, two-way ANOVA found no main effect of locomotion (F(1,19)=0.69, P=0.42) but revealed a significant locomotion x frequency interaction (F(1,19)=4.97, P=0.038); however, post-hoc testing found no difference between movement and freezing at low or high gamma frequencies.

## Discussion

Extensive evidence indicates that CFC is mediated by a neural circuit that includes the hippocampus, prefrontal cortex, and amygdala. However, the neurophysiological mechanisms underpinning the regulation of CFC by neuromodulators such as dopamine remain poorly understood. In this study we determined the effects of systemic D1R blockade on CFC and associated synchronized oscillations in the hippocampal-prefrontal-amygdala circuit. The selective D1R antagonist SCH23390 impaired CFC and acted acutely to alter intra-hippocampal and reduce hippocampal-prefrontal and hippocampal-amygdala synchrony during conditioning. SCH23390 decreased peak theta coherence and increased gamma coherence and theta-gamma PAC between DH and VH, while VH-PL and VH-BLA theta and gamma coherence were decreased. Impaired CFC by SCH23390 was associated with altered intra-hippocampal and reduced hippocampal-prefrontal-amygdala synchrony at retrieval. Prior SCH23390 decreased peak low gamma coherence and increased theta coherence and theta-gamma PAC between DH and VH, while resulting in decreased VH-PL, VH-BLA, and PL-BLA theta and gamma coherence (Fig 8). This reduction in hippocampal-prefrontal-amygdala synchrony was not fully accounted for by non-specific increases in movement occurring with decreased freezing at retrieval.

**Fig 8.**
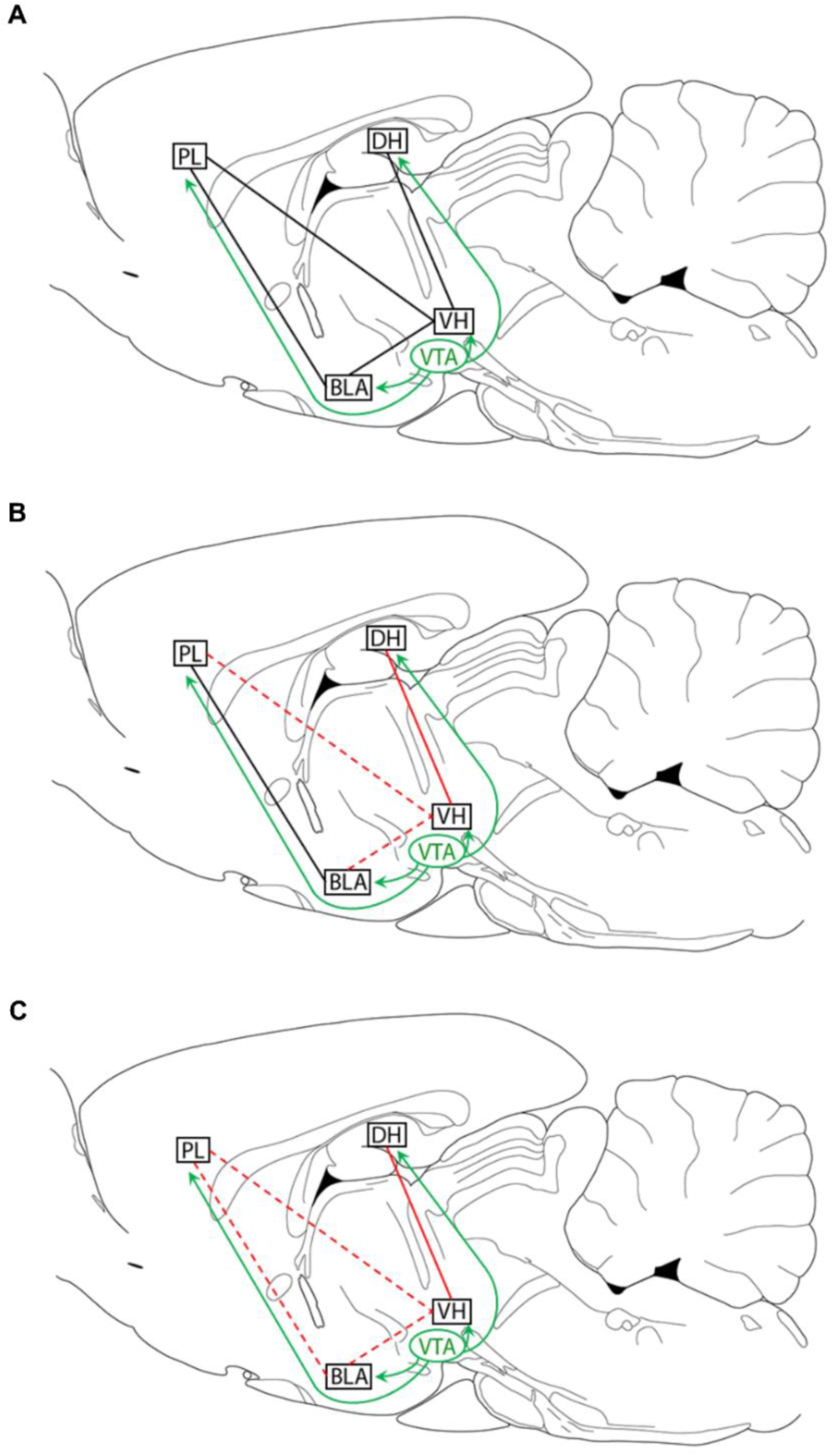
Summary of the key effects of D1R blockade before CFC on hippocampal-prefrontal-amygdala synchrony during conditioning acutely and at retrieval tested drug-free. A) Dopamine projections from the ventral tegmental area (VTA) to DH, PL, and BLA (green lines) regulate CFC and local synaptic plasticity via D1R signalling. DH projects to PL and BLA indirectly through VH (black lines). B) SCH23390 acts acutely during conditioning to alter intra-hippocampal (solid red line) and reduce hippocampal-prefrontal and hippocampal-amygdala (dashed red lines) theta and gamma synchrony. C) Impaired CFC by SCH23390 is associated with altered intra- hippocampal (solid red line) and reduced hippocampal-prefrontal-amygdala (dashed red lines) theta and gamma synchrony at retrieval.

This was revealed by comparing between epochs of movement and freezing in the vehicle-treated controls, which showed that movement increased VH-BLA and PL-BLA theta coherence and had no effect on VH-BLA or PL-BLA gamma coherence (Table 1). These results provide evidence that D1R signalling regulates CFC by modulating synchronized oscillations in the hippocampal-prefrontal- amygdala circuit, which also reflect a neural signature of contextual fear memory.

**Table 1.** Comparison of effects of impaired CFC by SCH23390 vs movement in the vehicle-treated controls on hippocampal-prefrontal-amygdala synchrony at retrieval.

| Areas | Synchrony measure | SCH23390 | Movement |
| --- | --- | --- | --- |
| DH-VH | theta coherence | ↑ | ↑ |
|  | gamma coherence | ↓/↑ | ↓ |
|  | theta-gamma PAC | ↑ | ↔ |
| VH-PL | theta coherence | ↓ | ↓ |
|  | gamma coherence | ↓ | ↓ |
|  | theta-gamma PAC | ↔ | ↔ |
| VH-BLA | theta coherence | ↓ | ↑ |
|  | gamma coherence | ↓ | ↔ |
|  | theta-gamma PAC | ↔ | ↔ |
| PL-BLA | theta coherence | ↓ | ↑ |
|  | gamma coherence | ↓ | ↔ |
|  | theta-gamma PAC | ↔ | ↔ |

Our finding that SCH23390 impaired CFC, as indicated by decreased freezing at retrieval in comparison to vehicle, replicates our previous results [Heath et al., 2015; Stubbendorff et al., 2019a]. In those studies, we showed that SCH23390 impairs the acquisition, but not consolidation, of contextual fear since SCH23390 given before, but not immediately after, conditioning reduced freezing at retrieval. Moreover, this effect of SCH23390 did not involve state dependency, acute drug effects on shock sensitivity during conditioning, or lasting drug effects on locomotion at retrieval. This indicates that D1R signalling at the time of conditioning is important for contextual fear learning.

Systemic or local D1R blockade has acute effects on hippocampal [Bo & Savoldi, 1992; Weiss et al., 2003; Xu et al., 2016; Park et al., 2021; Gonzalez et al., 2021], prefrontal [Parker et al., 2014; Ott et al., 2018], and amygdala [Loretan et al., 2004] oscillations. However, less is known about D1R modulation of synchronized oscillations between these inter-connected areas. Intra-cerebroventricular SCH23390 infusion reduces theta phase synchrony between VH and PL under anesthesia [Xu et al., 2016]. SCH23390 infused into VH prevents the increase in phase-locking of PFC spike firing to VH theta that occurs during learning [Park et al., 2021]. Our results showing that SCH23390 decreased theta and gamma coherence between VH and PL during CFC broadly agrees with these previous findings. We also found that SCH23390 decreased theta and gamma coherence between VH and BLA during CFC without affecting PL-BLA synchrony. In contrast, the effects of SCH23390 on DH-VH synchrony were more complex. SCH23390 decreased peak theta coherence but increased gamma coherence and theta-gamma PAC. This suggests that D1R signalling has different effects on the modulation of intra- hippocampal coupling and synchrony between hippocampus and its projection areas.

Our finding that impaired CFC by SCH23390 was characterized by reduced hippocampal-prefrontal-amygdala synchrony at retrieval generally agrees with evidence indicating that synchronized oscillations in this circuit are crucial for fear memory. Theta synchrony between hippocampus and PFC [Miller et al., 2017] or amygdala [Seidenbecher et al., 2003; Pape et al., 2005; Albrecht et al., 2010] is enhanced during contextual fear retrieval. Moreover, hippocampal-amygdala theta synchrony is associated with long-term (24 h) but not short-term (30 min) contextual fear memory or fear expression *per se* [Pape et al., 2005]. We examined VH-PL and VH-BLA synchrony since hippocampal projections to these areas occur through VH [Pitkanen et al., 2000; Cenquizca & Swanson, 2007; Kim & Cho, 2017]. Our results showing that impaired CFC is associated with reduced VH-PL and VH-BLA theta coherence at retrieval confirm and extend these previous findings implicating hippocampal-prefrontal and hippocampal-amygdala theta synchrony in contextual fear memory. Our finding of reduced PL-BLA theta coherence with impaired CFC also broadly agrees with studies showing that prefrontal-amygdala theta synchrony is involved in auditory fear retrieval [Lesting et al., 2011, 2013]. Similarly, we found reduced VH-PL, VH-BLA, and PL-BLA gamma coherence at retrieval with impaired CFC, suggesting that gamma synchrony between these areas also plays a role in contextual fear retrieval. This is compatible with evidence indicating that BLA gamma oscillations are important for contextual memory consolidation and that gamma coherence between BLA and various inter-connected areas is enhanced during learning [Bauer & Paré, 2007; Popescu et al., 2009; Kanta et al., 2019]. However, impaired CFC was associated with more complex interactions between DH and VH at retrieval. While peak low gamma coherence between DH and VH was decreased, theta and higher gamma coherence and theta-gamma PAC were increased. This suggests that impaired CFC is characterized by differences in intra-hippocampal coupling and hippocampal-prefrontal-amygdala synchrony at retrieval.

A potential confounding issue for interpreting the effects of systemic D1R blockade on hippocampal-prefrontal-amygdala synchrony is that non-specific locomotor effects might be involved. SCH23390 acts acutely to decrease locomotion [Inoue et al., 2000; Heath et al., 2015], while decreased freezing at retrieval resulting from impaired CFC by SCH23390 has the opposite effect to increase movement. Therefore opposing effects of SCH23390 acutely and during later retrieval might be expected if non-specific locomotor effects are involved. Although SCH23390 had the opposite effects on peak theta and low gamma coherence between DH and VH acutely and at retrieval, there were similar effects on DH-VH theta-gamma PAC, VH-PL theta and gamma coherence, and VH-BLA theta and gamma coherence under both conditions. To address this issue, we also compared hippocampal-prefrontal-amygdala synchrony between epochs of movement and freezing at retrieval in the vehicle-treated controls. If reduced freezing and VH-PL-BLA synchrony with impaired CFC by SCH23390 reflected a non-specific increase in locomotion at retrieval, then we expected to find a similar synchrony profile with movement in the controls, whereas differences between these synchrony profiles may indicate specific effects on contextual fear memory. Our results showed both similarities and differences in these synchrony profiles (Table 1). Movement increased theta and decreased peak low gamma coherence between DH and VH in controls, as we found with SCH23390. Movement also decreased VH-PL theta and gamma coherence in controls, albeit to a lesser extent than with SCH23390. However, VH-BLA and PL-BLA theta coherence were increased with movement in controls but decreased with SCH23390. Movement also had no effects on VH-BLA or PL-BLA gamma coherence in controls, whereas SCH23990 resulted in decreased gamma coherence between these areas. These results suggest that while the effects of SCH23390 on DH-VH and VH-PL synchrony at retrieval may have involved non-specific increases in movement, BLA synchrony with VH and PL during retrieval was instead more likely to have reflected a neural signature of contextual fear memory.

In summary, this study lends support to the idea that D1Rs regulate CFC by modulating hippocampal-prefrontal-amygdala synchrony. Synaptic plasticity in this circuitry, which is disrupted by local D1R blockade [Huang & Kandel, 1995; Gurden et al. 2000; Li et al., 2011], is facilitated by synchronized oscillations via their entrainment of correlated firing in neuronal populations [Headley & Paré, 2013; Bocchio et al., 2017; Totty & Maren, 2022]. Reduced synchrony between these areas is therefore a plausible neurophysiological mechanism by which D1R blockade disrupts the synaptic plasticity underlying CFC. This has implications for understanding how the various psychological processes involved are mediated by communication between these areas. D1R blockade locally in DH impairs CFC and LTP in this area [Huang & Kandel, 1995; Heath et al., 2015], which may interfere with encoding the context representation and context-US association. Although blocking D1Rs locally in VH does not affect CFC [Stubbendorff et al., 2019a], our results suggest that altered synchrony between DH and VH by systemic D1R blockade may also play a role in disrupting CFC. Moreover, reduced VH-PL and VH-BLA synchrony with systemic D1R blockade may impair CFC by interfering with the context and US representations being conveyed from hippocampus to these areas [Kim & Cho, 2017; Jimenez et al., 2020]. In support of this idea, blocking D1Rs locally in PL impairs CFC and LTP in the VH-PL pathway [Gurden et al., 2000; Stubbendorff et al., 2019a]. Local D1R blockade in BLA also impairs CFC and LTP in this area [Li et al., 2011; Heath et al., 2015], which may interfere with encoding the US representation and context-US association [Tang et al., 2020]. However, D1R modulation of VH-BLA plasticity [Maren & Fanselow, 1995; Kim & Cho, 2020] remains to be determined. Further research is also needed to determine the source and cell type-specific targets of the dopamine projections that modulate synchrony in this circuit, along with the underlying local neuronal population dynamics. Nevertheless, this study builds on our understanding of the hippocampal-prefrontal-amygdala circuit mechanisms underlying the neuromodulation of CFC by dopamine. It also confirms and adds to evidence indicating that communication within this circuitry is crucial for fear memory [Headley & Paré, 2013; Bocchio et al., 2017; Totty & Maren, 2022] by characterizing synchrony within hippocampus and between prefrontal cortex, amygdala and ventral hippocampus, the source of hippocampal projections to these areas. This may ultimately lead to novel insights on the neurobiological basis of anxiety-related disorders characterized by aberrant learned fear processing.

## Methods

### Animals

42 male Lister Hooded rats (Charles River, UK) weighing 280-390 g before surgery were used. Rats were group housed in individually ventilated cages (2-3/cage) and kept on a 24 h light/dark cycle (lights on at 7:00) with *ad libitum* access to food and water. All behavioral testing occurred during the rats’ light cycle. All experimental procedures were performed with institutional ethical approval and under the UK Animals (Scientific Procedures) Act 1986 (Home Office Project Licence 30/3230).

### Electrode implant surgery

Anesthesia was induced with ∼3% isoflurane in oxygen and analgesic (buprenorphine, 0.05 mg/kg, s.c.) was administered immediately post-induction. Anesthesia was maintained with 1.5-2.5% isoflurane during surgery to ensure complete inhibition of the hindpaw withdrawal reflex. Rats were placed in a stereotaxic frame and the incisor bar was adjusted to maintain the skull horizontal. A homoeothermic heating pad was used to maintain body temperature at 36-37 °C throughout surgery. A scalp incision was made along the midline, the periosteum was retracted, and 10 stainless steel anchoring screws were affixed to the top and sides of the skull. Small craniotomies were performed on the right side above the target coordinates and the dura mater was incised immediately before electrode implantation. Custom-made tungsten electrodes [Stubbendorff et al., 2019b] were targeted at DH (CA1; 3.0 mm posterior and 1.5 mm lateral to bregma, 3.0 mm ventral to the brain surface), VH (CA1; 5.0 mm posterior and 4.8 mm lateral to bregma, 6.3 mm ventral to the brain surface), PL (2.5 mm anterior and 0.5-0.8 mm lateral to bregma, 3.0 mm ventral to the brain surface), and BLA (basal nucleus; 2.8 mm posterior and 4.7 mm lateral to bregma, 7.2 mm ventral to the brain surface) (Paxinos & Watson, 2007). The electrodes were loaded into a microdrive (VersaDrive-8, Neuralynx, MT) and the implant was secured to the anchoring screws with light-cured dental cement (Henry Schein, UK). Another analgesic (meloxicam, 1 mg/kg, s.c.) was given at the end of surgery. Rats were singly housed for 1-2 days after surgery, after which they were group housed as above. Buprenorphine and meloxicam were given once daily for 2-3 days after surgery. Behavioral testing commenced 6-8 days after surgery.

### Drug injection

SCH23390 hydrochloride (0.1 mg/kg, i.p.; Tocris Bioscience, UK) was dissolved in 0.9% sterile saline (1 mL/kg) and injected 30 min before CFC (see below). We have previously shown that this dose impairs CFC [Heath et al., 2015; Stubbendorff et al., 2019a]. Vehicle-treated controls received injections of 0.9% sterile saline (1 mL/kg, i.p.).

### Behavioral testing

The effects of systemic SCH23390 administration on CFC were investigated using a two-day conditioning and retrieval paradigm. Each rat was randomly allocated to receive SCH23390 or vehicle treatment before CFC. The apparatus [Stevenson et al., 2009] and experimental procedures [Stubbendorff et al., 2019a] used have been described in detail elsewhere. On the first day rats underwent conditioning in a novel context that consisted of distinct visual, auditory, and olfactory cues present in the background during CFC. The US was a mild electric shock delivered through the chamber floor bars automatically via a PC running Med-PC IV software (Med Associates, VT). The rat was placed in the chamber and after 2 min was presented with four shocks (0.5 mA, 0.5 s, 1 min inter-trial interval). The rat was removed from the chamber 2 min after presentation of the last shock and returned to the home cage. On the second day the rat was returned to the conditioning chamber for 5 min to test memory retrieval drug-free. The floor bars and waste tray were cleaned with 40% ethanol between each session. Rats were tested at approximately the same time of day on both days. Electrophysiological recordings (see below) were obtained during CFC and retrieval testing. Behavior during retrieval testing was recorded using a digital camera (ViewPoint, France) positioned above the chamber.

### Electrophysiological recordings

LFPs from the electrodes targeting each area were recorded during CFC and retrieval testing by connecting the microdrive via a headstage, cable, and pre-amplifier to an OmniPlex neural recording data acquisition system (Plexon Inc, TX). LFPs were band-pass filtered at 0.7-170 Hz and digitized at 1.25 kHz. A cable connecting the Med Associates and OmniPlex systems was used to record the start of the CFC and retrieval sessions, triggered by the Med-PC IV software, in the LFP data recording file.

### Histology

After completing retrieval testing, rats were deeply anesthetized with sodium pentobarbital and current was passed through each electrode using an electrical stimulator to create a small lesion at the electrode tips. Rats were then perfused transcardially with 0.9% saline followed by 4% paraformaldehyde. The brains were removed, post-fixed in 4% paraformaldehyde, and kept at 4 °C until slicing. Sections containing the relevant areas were obtained using a vibratome and stained for acetylcholinesterase (Fig 1D). Only data from rats with histologically confirmed electrode placements in DH, VH, PL, and/or BLA were included in the electrophysiological data analysis.

### Behavioral data analysis

Freezing, defined as the absence of movement except in relation to respiration, was taken as the behavioral measure of contextual fear during retrieval testing. Freezing was quantified automatically using VideoTrack software (ViewPoint, France) by setting the freezing detection threshold to 100 pixels change/frame (25 frames/sec), based on a comparison between automatically determined and manually scored freezing levels from our published data [Stubbendorff et al., 2019a]. The cumulative duration of freezing during retrieval testing was calculated and expressed as a percentage of the 5 min test duration. Differences in freezing between the two groups were analyzed in two ways. Average freezing over the whole 5 min test session was analyzed using a two-tailed unpaired t-test, with the data presented in a bar graph as the mean + standard error of the mean (SEM). Freezing during each 1-min bin was also analyzed separately using two-way analysis of variance (ANOVA), with treatment as the between-subjects factor and time as the within-subjects factor. Freezing during each 1-min bin was presented in a line graph as the mean + SEM. The level of significance for both analyses was set at P<0.05.

### Electrophysiological data analysis

LFP activity was analyzed using multi-taper spectral analysis as described elsewhere [Fenton et al., 2013, 2014; Day et al., 2020]. LFP data were high-pass filtered at 4 Hz and inspected visually, with LFP recordings containing obvious electrical noise artefacts omitted from the analysis. LFP signals were normalized to unit variance in each segment prior to spectral analysis. Normalization removes effects of changes in absolute power due to changes in electrode impedance, therefore we report on differences in relative power. Spectral estimates for LFP data from each area in the 2 min period before US presentations during CFC and in the 5 min retrieval test were generated using custom Matlab scripts. All electrophysiology data and data analysis code are freely available online (https://osf.io/km2s7/).

We first examined the acute effects of SCH23390 on theta (4-12 Hz), low gamma (30-45 Hz), and high gamma (55-80 Hz) power in each area during CFC. The gamma frequency band was divided into low and high frequencies to omit electrical mains noise ∼50 Hz. Differences between SCH23390 and vehicle treatment in power at each individual frequency throughout the theta and gamma bands were quantified using the log ratio test [Diggle, 1990]. The acute effects of SCH23390 on theta, low gamma, and high gamma coherence between the areas sharing direct anatomical connections (i.e. DH-VH, VH-PL, VH-BLA, and PL-BLA) were then determined during CFC as measures of synchrony between these areas. Differences between SCH23390 and vehicle treatment in coherence at each individual frequency throughout the theta and gamma bands were quantified using the chi-squared difference of coherence test [Amjad et al., 1997]. Power and coherence data are presented as the mean +/- 95% pointwise confidence intervals. The level of significance for the statistical comparisons of power and coherence was set at P<0.001 to correct for multiple comparisons across individual frequencies. Differences at individual frequencies were also considered as chance effects, therefore differences were only deemed significant where they were found for two or more adjacent frequencies. The acute effects of SCH23390 on theta-gamma PAC during CFC were quantified as described elsewhere [Canolty et al., 2006; Samiee & Baillet, 2017; Suwansawang & Halliday, 2017] using custom Matlab scripts. Theta-gamma PAC was determined using a modulation index from the normalised mean vector length. Estimates were constructed using analytic Morse wavelets, with significance determined using surrogate data [Lilly & Olhede, 2012; Aru et al., 2015; Suwansawang & Halliday, 2017]. We examined theta-gamma PAC within each area and between the directly inter-connected areas. Specifically, we examined DH theta-VH gamma PAC since theta oscillations propagate from DH to VH [Lubenov & Siapas, 2009; Patel et al., 2012]. We also examined VH theta-PL gamma PAC since hippocampal-prefrontal theta synchrony is mediated by VH [Adhikari et al., 2010; O’Neill et al., 2013]. Similarly, we examined VH theta-BLA gamma PAC since theta burst stimulation of VH modulates BLA activity and plasticity [Bazelot et al., 2015]. Finally, we examined PL theta-BLA gamma PAC since learned fear inhibition is associated with prefrontal theta coupling of BLA gamma [Stujenske et al., 2014]. Color plots of theta-gamma PAC within each area and between the directly inter-connected areas showed that peak theta phase coupling of gamma amplitude occurred at a theta frequency ∼5 Hz after vehicle or SCH23390 treatment, therefore we focused the analyses at this theta frequency. Theta-gamma PAC was analyzed using two-way ANOVA, with treatment as the between-subjects factor and gamma frequency band as the within-subjects factor. Post-hoc comparisons were conducted using the Sidak test where indicated. Theta-gamma PAC was presented in bar graphs as the mean + SEM and the level of significance for the analyses was set at P<0.05.

The effects of SCH23390 given before CFC on theta power and coherence, low and high gamma power and coherence, and theta-gamma PAC during retrieval testing drug-free were determined as above. We also examined differences in these neurophysiological measures between epochs of movement and freezing during retrieval testing in the vehicle-treated controls to determine if decreased freezing resulting from SCH23390 treatment before CFC was related specifically to impaired contextual fear or non-specifically to increased movement at retrieval. Freezing epochs were determined by considering intervals of a minimum of 0.5 sec where <100 pixels change/frame (i.e. the freezing detection threshold) was detected, while movement epochs were determined by considering intervals of a minimum of 0.5 sec where >500 pixels change/frame was detected. The resulting LFP data from these intervals were then pooled within and between rats. Differences between movement and freezing in theta power and coherence, low and high gamma power and coherence, and theta-gamma PAC during retrieval testing were determined as above.

## Acknowledgements

This work was funded by a research grant from the Biotechnology and Biological Sciences Research Council [grant number BB/P001149/1].

## Supplementary Tables

**Table S1.** Statistical analysis of acute SCH23390 effects on theta-gamma PAC during conditioning.

| **Within area** |  | **F value** | **P value** |
| --- | --- | --- | --- |
| DH-DH | Main effect of treatment | $F_{(1,31)}=0.88$ | $P=0.36$ |
| | Main effect of frequency | $F_{(1,31)}=65.48$ | $P<0.0001$ |
| | Treatment x frequency interaction | $F_{(1,31)}=0.84$ | $P=0.37$ |
| VH-VH | Main effect of treatment | $F_{(1,28)}=3.19$ | $P=0.085$ |
| | Main effect of frequency | $F_{(1,28)}=73.84$ | $P<0.0001$ |
| | Treatment x frequency interaction | $F_{(1,28)}=2.05$ | $P=0.16$ |
| PL-PL | Main effect of treatment | $F_{(1,26)}=0.23$ | $P=0.64$ |
| | Main effect of frequency | $F_{(1,26)}=106.0$ | $P<0.0001$ |
| | Treatment x frequency interaction | $F_{(1,26)}=1.28$ | $P=0.27$ |
| BLA-BLA | Main effect of treatment | $F_{(1,16)}=1.45$ | $P=0.25$ |
| | Main effect of frequency | $F_{(1,16)}=76.26$ | $P<0.0001$ |
| | Treatment x frequency interaction | $F_{(1,16)}=1.22$ | $P=0.29$ |

| **Between areas** |  | **F value** | **P value** |
| --- | --- | --- | --- |
| DH theta-VH gamma | Main effect of treatment | $F_{(1,24)}=4.41$ | $P=0.046$ |
| | Main effect of frequency | $F_{(1,24)}=74.92$ | $P<0.0001$ |
| | Treatment x frequency interaction | $F_{(1,24)}=1.75$ | $P=0.20$ |
| VH theta-PL gamma | Main effect of treatment | $F_{(1,21)}=1.61$ | $P=0.22$ |
| | Main effect of frequency | $F_{(1,21)}=120.8$ | $P<0.0001$ |
| | Treatment x frequency interaction | $F_{(1,21)}=4.53$ | $P=0.045$ |
| VH theta-BLA gamma | Main effect of treatment | $F_{(1,12)}=0.27$ | $P=0.61$ |
| | Main effect of frequency | $F_{(1,12)}=51.42$ | $P<0.0001$ |
| | Treatment x frequency interaction | $F_{(1,12)}=0.16$ | $P=0.69$ |
| PL theta-BLA gamma | Main effect of treatment | $F_{(1,12)}=0.16$ | $P=0.70$ |
| | Main effect of frequency | $F_{(1,12)}=61.61$ | $P<0.0001$ |
| | Treatment x frequency interaction | $F_{(1,12)}=0.028$ | $P=0.87$ |

**Table S2.** Statistical analysis of SCH23390 effects on theta-gamma PAC at retrieval.

| **Within area** |  | **F value** | **P value** |
| --- | --- | --- | --- |
| DH-DH | Main effect of treatment | $F_{(1,35)}=0.17$ | $P=0.69$ |
| | Main effect of frequency | $F_{(1,35)}=64.96$ | $P<0.0001$ |
| | Treatment x frequency interaction | $F_{(1,35)}=0.34$ | $P=0.56$ |
| VH-VH | Main effect of treatment | $F_{(1,30)}=3.08$ | $P=0.089$ |
| | Main effect of frequency | $F_{(1,30)}=37.85$ | $P<0.0001$ |
| | Treatment x frequency interaction | $F_{(1,30)}=0.82$ | $P=0.37$ |
| PL-PL | Main effect of treatment | $F_{(1,28)}=0.45$ | $P=0.51$ |
| | Main effect of frequency | $F_{(1,28)}=6.30$ | $P=0.018$ |
| | Treatment x frequency interaction | $F_{(1,28)}=1.49$ | $P=0.23$ |
| BLA-BLA | Main effect of treatment | $F_{(1,17)}=1.62$ | $P=0.22$ |
| | Main effect of frequency | $F_{(1,17)}=18.66$ | $P=0.0005$ |
| | Treatment x frequency interaction | $F_{(1,17)}=0.034$ | $P=0.86$ |

| **Between areas** |  | **F value** | **P value** |
| --- | --- | --- | --- |
| DH theta-VH gamma | Main effect of treatment | $F_{(1,27)}=5.03$ | $P=0.03$ |
| | Main effect of frequency | $F_{(1,27)}=33.87$ | $P<0.0001$ |
| | Treatment x frequency interaction | $F_{(1,27)}=1.70$ | $P=0.20$ |
| VH theta-PL gamma | Main effect of treatment | $F_{(1,23)}=0.79$ | $P=0.38$ |
| | Main effect of frequency | $F_{(1,23)}=6.75$ | $P=0.016$ |
| | Treatment x frequency interaction | $F_{(1,23)}=0.66$ | $P=0.42$ |
| VH theta-BLA gamma | Main effect of treatment | $F_{(1,14)}=1.28$ | $P=0.28$ |
| | Main effect of frequency | $F_{(1,14)}=20.40$ | $P=0.0005$ |
| | Treatment x frequency interaction | $F_{(1,14)}=0.71$ | $P=0.41$ |
| PL theta-BLA gamma | Main effect of treatment | $F_{(1,13)}=1.64$ | $P=0.22$ |
| | Main effect of frequency | $F_{(1,13)}=22.39$ | $P=0.0004$ |
| | Treatment x frequency interaction | $F_{(1,13)}=0.85$ | $P=0.37$ |

**Table S3.** Statistical analysis of movement effects on theta-gamma PAC at retrieval in the controls.

| **Within area** |  | **F value** | **P value** |
| --- | --- | --- | --- |
| DH-DH | Main effect of locomotion | $F_{(1,25)}=0.040$ | P=0.84 |
| | Main effect of frequency | $F_{(1,25)}=33.68$ | P<0.0001 |
| | Locomotion x frequency interaction | $F_{(1,25)}=0.58$ | P=0.45 |
| VH-VH | Main effect of locomotion | $F_{(1,21)}=0.10$ | P=0.75 |
| | Main effect of frequency | $F_{(1,21)}=12.83$ | P=0.0018 |
| | Locomotion x frequency interaction | $F_{(1,21)}=0.34$ | P=0.57 |
| PL-PL | Main effect of locomotion | $F_{(1,16)}=1.04$ | P=0.32 |
| | Main effect of frequency | $F_{(1,16)}=10.43$ | P=0.0052 |
| | Locomotion x frequency interaction | $F_{(1,16)}=1.13$ | P=0.30 |
| BLA-BLA | Main effect of locomotion | $F_{(1,7)}=4.05$ | P=0.084 |
| | Main effect of frequency | $F_{(1,7)}=23.63$ | P=0.0018 |
| | Locomotion x frequency interaction | $F_{(1,7)}=5.59$ | P=0.050 |

| **Between areas** |  | **F value** | **P value** |
| --- | --- | --- | --- |
| DH theta-VH gamma | Main effect of locomotion | $F_{(1,19)}=0.69$ | P=0.42 |
| | Main effect of frequency | $F_{(1,19)}=13.22$ | P=0.0018 |
| | Locomotion x frequency interaction | $F_{(1,19)}=4.97$ | P=0.038 |
| VH theta-PL gamma | Main effect of locomotion | $F_{(1,11)}=0.93$ | P=0.36 |
| | Main effect of frequency | $F_{(1,11)}=4.97$ | P=0.048 |
| | Locomotion x frequency interaction | $F_{(1,11)}=0.86$ | P=0.37 |
| VH theta-BLA gamma | Main effect of locomotion | $F_{(1,6)}=1.66$ | P=0.24 |
| | Main effect of frequency | $F_{(1,6)}=18.21$ | P=0.0053 |
| | Locomotion x frequency interaction | $F_{(1,6)}=2.29$ | P=0.18 |
| PL theta-BLA gamma | Main effect of locomotion | $F_{(1,6)}=1.79$ | P=0.23 |
| | Main effect of frequency | $F_{(1,6)}=22.69$ | P=0.0031 |
| | Locomotion x frequency interaction | $F_{(1,6)}=1.84$ | P=0.22 |

